# Untangling the dynamics of persistence and colonization in microbial communities

**DOI:** 10.1101/427542

**Authors:** Sylvia L. Ranjeva, Joseph R. Mihaljevic, Maxwell B. Joseph, Anna R. Giuliano, Greg Dwyer

## Abstract

A central goal of community ecology is to infer biotic interactions from observed distributions of co-occurring species. Evidence for biotic interactions, however, can be obscured by shared environmental requirements, posing a challenge for statistical inference. Here we introduce a dynamic statistical model that quantifies the effects of spatial and temporal covariance in longitudinal co-occurrence data. We separate the fixed pairwise effects of species occurrences on persistence and colonization rates, a potential signal of direct interactions, from latent pairwise correlations in occurrence, a potential signal of shared environmental responses. We apply our modeling approach to a pressing epidemiological question by examining how human papillomavirus (HPV) types coexist. Our results suggest that while HPV types respond similarly to common host traits, direct interactions are sparse and weak, so that HPV type diversity depends largely on shared environmental drivers. Our modeling approach is widely applicable to microbial communities and provides valuable insights that should lead to more directed hypothesis testing and mechanistic modeling.

## Introduction

A fundamental goal of community ecology is to understand how interactions between species in a shared environment shape observed patterns of diversity over time. A key challenge in understanding community turnover is to disentangle effects of environmental drivers of species co-occurrence from inter-species interactions, especially when the goal is to infer these mechanisms from observational data [1, 2]. This challenge is also found in epidemiology, in which a major goal is to understand the factors that allow pathogens to coexist [3]. As is the case with free-living species, when determinants of environmental niches are shared among pathogen types, inferring interactions is difficult [4]. Understanding the mechanisms of microbial community turnover thus presents an ecological, statistical, and computational challenge, especially considering the size of microbial and pathogen data sets [5, 6]. Ecological models of community turnover that account for shared environmental drivers are thus important for understanding mechanisms that underlie pathogen diversity.

For macroscopic organisms, null model analysis has historically been used to infer potential species interactions from observational data sets, through the identification of statistically non-random aggregations of species across multiple habitats [1, 7–9]. Similar approaches have been used to develop computationally efficient algorithms that make it possible to infer large correlation networks from microbial sequence data [5, 10]. Disentangling the simultaneous effects of species interactions and environmental filters from survey data is nevertheless a challenge for analyses of both macroscopic and microscopic communities [2, 11]. For example, highly mobile, competing species should transiently aggregate in habitats with high shared resources, even if competitive exclusion is expected at equilibrium. Snap-shot surveys of co-occurrence can therefore lead to biased interpretations of species interactions, but time-series data can help overcome this problem.

In the microbial ecology literature, network inference models have only rarely been adapted to incorporate time-series data from multiple localities. Available methods include local similarity analysis [11–13] and generalized Lotka-Volterra modeling [14, 15]. While local similarity analysis can be used with incidence data, Lotka-Volterra modeling requires measures of abundance, which are notoriously difficult to infer from sequence data, whereas relative abundances can bias statistical analyses [16]. Local similarity analysis can infer microbial networks from observations of time-delays and temporal correlations between microbes and environmental covariates, but it relies on multiple, independent tests with p-value corrections, instead of an integrated analysis [12, 13]. In addition, simulation studies have determined that correlations (e.g., Pearson’s correlations or time-lagged correlations) between species’ raw abundances are often not indicative of underlying species interactions [10, 17].

Joint species distribution models provide a more comprehensive method for identifying putatively interacting species from static ecological survey data, while accounting for shared environmental drivers [18–23]. These models use logistic regression to estimate how environmental covariates affect species occupancy probabilities across a heterogeneous landscape. Species interactions are then inferred from residual correlations between species occurrences. Although these residual correlations are better accounting for environmental effects, their interpretation as species interactions is still problematic, and should be used only for hypothesis generation. Furthermore, while joint-species models can generate hypotheses about static community assemblages, most methods fail to capture important drivers of co-occurrence that emerge from dynamic properties of the community dynamics [2]. For example, species co-occurrence may be positively correlated across heterogeneous habitats, because of shared resources, but negatively correlated across time, because of negative species interactions within sites. This is an example of Simpson’s paradox, in which positive associations at one scale can obscure negative associations at a smaller scale. And, this pattern is a ubiquitous problem for ecological studies (Fig. 1).

**Figure 1:**
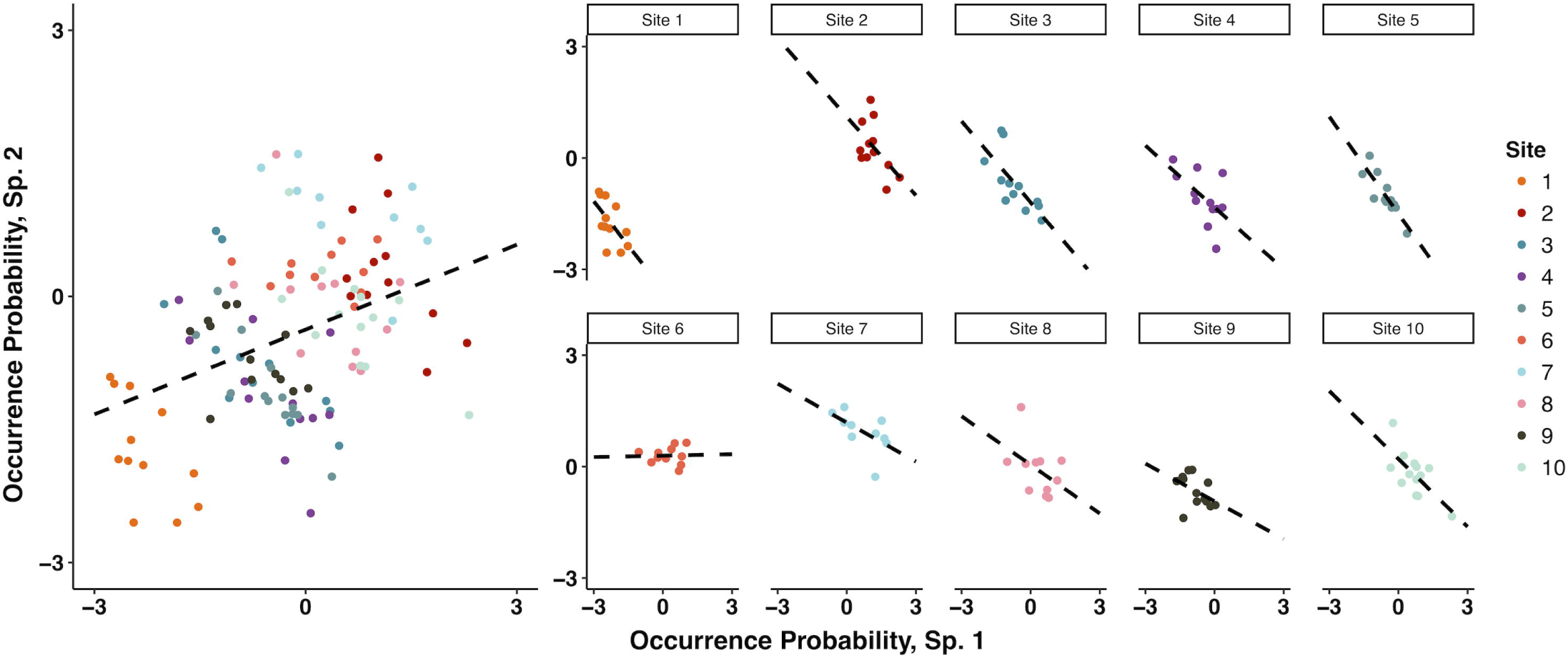
Simpson’s paradox demonstrated for two species that are sampled across ten habitat sites, with each site surveyed fifteen times. **A** Species covary positively across sites (over space), indicating response to similar habitat requirements. **B** Species covary negatively within sites over time, indicating inter-specific competition. Probabilities of occurrence are on the probit scale.

Here we extend the joint-species modeling framework to infer more complex, biologically realistic dynamics in a way that is computationally tractable for large microbial data sets. We develop a statistical model of a dynamic, multi-species metacommunity in which species’ occurrences can affect other species’ persistence and colonization probabilities (fixed effects), and occurrences are further influenced by shared environmental drivers (random effects). This approach can be readily applied to pathogenic microbe populations, in which distinct pathogen types represent species coexisting within a heterogeneous landscape of host organisms. In a generalized, linear mixed effects framework, we model correlations in species occupancy across habitats and across time by allowing for correlations in random effects at different spatial and temporal scales, resolving Simpson’s paradox and accounting for latent environmental covariates. We also estimate pairwise species effects on rates of colonization and persistence as fixed effects. Using synthetic data, we demonstrate the ability of our model to accurately and precisely infer dynamics consistent with Simpson’s paradox, even with sparse occurrences, and across a range of data set characteristics (e.g., number of patients sampled, and sizes of fixed and random effects). We then apply our model to data on human papillomavirus (HPV), a pathogen of significant public health concern.

Human papillomavirus (HPV) is the most common sexually transmitted infection and a major cause of cervical, genital, and oropharyngeal cancers, and it consists of over 200 types [24]. Uncertainty about the mechanisms underlying HPV type coexistence, and particularly about potential HPV type interactions, reflects a crucial unknown. Four HPV types cause most disease symptoms [24–26] and quadrivalent vaccination has demonstrated high efficacy in reducing rates of cervical dysplasia and genital warts [27, 28]. A recent 9-valent HPV vaccine targets additional oncogenic types [29]. Because the HPV vaccine is multivalent, it is possible that type replacement will occur, in which non-vaccine types increase in frequency due to population-level removal of vaccine-targeted types [30]. Type replacement following vaccination depends on interactions between HPV types during natural infection, and particularly on inter-type competition through cross-immunity [31]. Understanding the ecological mechanisms that underlie HPV type diversity could therefore inform strategies for disease management and prevention. It has thus far been difficult to distinguish HPV type interactions from the effects of shared host-specific risk factors. Our dynamical community model allows us to investigate how type interactions and risk factors together structure the HPV viral community.

In this study we address two questions, which differ in their scope. First, we use our full model to ask which interactions between specific HPV types warrant future investigation? Second, we ask a more ecological question: what are the dominant drivers of community composition across space and time? To address this second question, we build models of increasing complexity, and we use model selection to determine whether HPV community patterns are determined by putative interactions between HPV types, by host-level factors that determine HPV distributions, or both. Our full model identified several interactions that warrant further experimental investigation, including negative pairwise effects on persistence and colonization probabilities. In addition, there is a strong signal of shared environmental drivers among HPV types, highlighting the importance of host-specific risk factors in supporting coexistence. By comparing models of varying complexity, however, we show that the dynamics of the HPV community are most parsimoniously explained by shared environmental drivers, rather than by strong pairwise interactions between HPV types. Pairwise species interactions thus do not appear to drive community-wide patterns of co-occurrence in the HPV community. Our study demonstrates the ability of our joint-species models to quickly and efficiently infer properties of a large, real-world viral community, and the model could therefore be of broad usefulness in understanding microbial communities.

## Materials and Methods

### HPV natural history

HPV types are classified based on the L1 viral capsid protein. A distinct HPV type is a variant whose L1 gene sequence is at least 10% dissimilar from any other HPV type [32]. The transmission and coexistence of individual HPV types depend on traits and risk factors of individual hosts [33–36]. These include determinants of sexual behavior, including frequency of condom use, number of new and steady sexual partners, and sexual orientation; demography, including race and ethnicity; and non-sexual behavior, including smoking and alcohol consumption.

Interactions between HPV types could determine HPV diversity, though conclusive evidence of HPV type interactions is lacking [31, 37, 38]. As in any species, HPV type interactions may be synergistic, neutral, or competitive. Synergism occurs when one type facilitates infection by another, while competition occurs when one type prevents infection by another. Under competitive interactions, removal of one HPV type should lead to an increase in prevalence of the competing type in the host population, resulting in type replacement. Natural history surveys reporting elevated odds ratios for multiple to single infections with HPV have suggested that cross-immunity among HPV types is unlikely [39–42]. Additionally, the genetic stability of HPV as a double-stranded DNA virus has been used to support arguments against the possibility of type replacement [43], on the grounds that rapid emergence of antigenic variants is unlikely [28]. Nevertheless, a recent increase in prevalence of non-vaccine types was found in young women following vaccination and in the United States [37], suggesting that type replacement may be occurring. Indeed, several models of HPV type interactions indicate that competition between HPV types is plausible under observed patterns of coinfections [31, 44] and have demonstrated the possibility of type-replacement after vaccination [31, 44–46].

### Data

We fit models of HPV type dynamics to data from the HPV Infection in Men (HIM) study [33, 34, 47], a multinational cohort study of HPV infection in men with no prior diagnosis of genital cancer or other sexually transmitted infections. The HIM study enrolled over 4000 men between 2005 and 2009 from three cities: Tampa, Florida, USA; Cuernavaca, Mexico; and Sao Paulo, Brazil. Detailed study methods are described elsewhere [33]. Briefly, the HIM study tracked PCR-confirmed infections with 37 types of HPV in men over a mean of 5 years of follow-up, recording behavioral and demographic information for all participants. The data for each individual consist of binary time series describing infection status with respect to each type over a median of 10 clinic visits, at median intervals of 6.0 months (variance = 0.7 months).

For the present analysis, we included the 3656 participants with no reported diagnosis of HIV and with PCR tests for each HPV type at all clinic visits (see Appendix). Our selection of HPV types was motivated by clinical and ecological relevance, HPV type prevalence, and computational tractability. We limited the analysis to ten of the HPV types represented in the HIM dataset: the nine HPV types included in the most recent HPV vaccine [29] and HPV84, a type that has shown high prevalence in several studies among men [24, 48]. We therefore accounted for seven high-risk types - HPV16, HPV18, HPV31, HPV33, HPV45, HPV52, and HPV58 – that together account for over 70% of HPV-associated malignancies in men and women, as well as three low-risk types - HPV6, HPV11, and HPV84 – that together are responsible for over 90% of benign anogenital lesions [49]. Additionally, ecological interactions between these vaccine-targeted types could lead to type replacement [31, 43, 45]. Finally, many of the other HPV types were quite rare: fewer than half of the 37 available HPV types have a mean prevalence greater than 2.5% over the course of the study. With these10 HPV types, our study includes 305,250 data points (one point per patient per virus type per visit), which is a computational burden, but manageable with our model.

### Statistical Model

Our goal is to extend current joint-species modeling techniques to biological processes that may be needed to understand community dynamics. Currently, only a limited number of joint-species modeling techniques are available for longitudinal survey data. Sebastian-Gonzalez et al. [21] extended the joint-species modeling framework to allow for multiple community surveys through time by modeling the fixed, pairwise effects of species co-occurrence between subsequent time points. Then, the model accounts for shared environmental drivers by essentially adding a random effect of habitat patch and allowing for pairwise correlations in these random effects, such that species can share similar responses to latent environmental variables. Dorazio et al. [50] introduced a model that separately estimated rates of species colonization and persistence from sequential community surveys. Although this latter model specifies the processes of extinction and colonization that can explain occupancy dynamics over time, it does not account for the residual dependence among species that can result from species interactions. Here we describe a statistical model that is tailored to the repeated surveys of patients in the HIM dataset, thereby combining the methods of Sebastian-Gonzalez et al. [21] and Dorazio et al. [50] in a computationally tractable and generalizable way.

Our data consist of observations made in *I* patients, who can harbor up to *J* HPV types (in our case limited to 10 types), sampled over a maximum of *T* sequential visits to the clinic. Observations of the HPV dataset are therefore aggregated as binary presence/absence data in the *I* × *J* × *T* incidence array **Y**, such that *Y*_*i,j,t*_ indicates the presence or absence of HPV type *j* in patient *i* at visit *t*.

We fit a multivariate probit regression model to the binary presence/absence data in **Y**, which has been used in other joint-species modeling approaches [22]. Probit regression relates a linear predictor to occupancy probabilities using a standard normal cumulative distribution function. In this model, the probability that a binary random variable is equal to one (i.e. *P*(*Y = 1*)) is equal to the probability that the latent variable *z* is greater than zero. The linear predictor *µ* completely determines the latent variable *z* and can be a function of one or more covariates and their effects. As part of the probit definition, the residual variance of *z* is equal to one. In general then, we are interested in understanding how linear predictors influence the probability that an HPV type occurs in a given patient. A generalized probit model with a single covariate *x* is formulated for the *i*^*th*^ sample as:

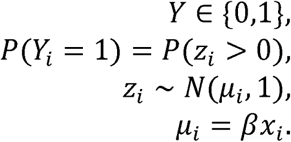

Our model extends the generalized probit model by assuming that occurrence probabilities are affected by both patient-level effects and potential interactions between HPV types. We therefore build upon the general case of the probit model to model observations of the dynamic HPV metacommunity. To account for temporal dynamics, we assume that the linear predictor *µ* _*i,j,t*_ for each observation depends on observation-specific probabilities of persistence and colonization:

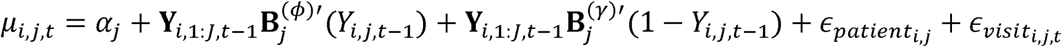

Here, *α*_*j*_ is an adjustment to account for among-type variation in commonness. The presence of a given HPV type can affect the probability of persistence or colonization of other types, with a one time-step lag. If HPV type *j* was present in patient *i* on the previous clinic visit (*t − 1*), then persistence effects are represented by the product 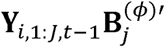, where *Y*_*i*,1:*J,t* −1_ is a row vector of length *J* containing the presence/absence states of strains *j =* 1,…,*J* in patient *i* on the previous visit (*t − 1*), and 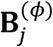 is a column vector of length *J* containing pairwise interaction coefficients (fixed effects). These coefficients thus specify how HPV type composition at the previous visit affects persistence (*ϕ*) of type *j*. We note that the special case of *β*_*jj*_ represents the effect of type *j* on its own persistence. In other words, if type *j* was present in visit *t, β*_*jj*_ affects the likelihood that this type will persist in a patient to the next visit.

If type *j* was absent in patient *i* on visit *t − 1*, colonization effects are represented by the product 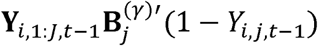, where 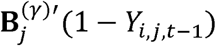 is a column vector of length *J*, again containing pairwise interaction coefficients (fixed effects). These coefficients thus specify how HPV type composition at the previous visit affects the colonization (*γ*) of type *j*. Both interaction matrices (*B*^(*ϕ*)^ and *B*^(*γ*)^) are *J*×*J* dimensional, and 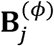 and 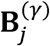 represent the row vectors acquired by extracting row *j*.

Lastly, patient-level and visit-level adjustments are specified as random effects 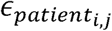 and 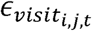 respectively. The multivariate patient-level random effect *∈*_*patient*_ allows pairwise correlations in HPV type occurrence across patients, thereby describing pairwise similarities in environmental requirements. In the case of the HIM data, *∈*_*patient*_ therefore controls for shared determinants of host risk, such as host behavioral covariates, that could confound estimates of HPV type interactions. The random visit-level effect *∈*_*visit*_ allows for pairwise correlations in HPV type occurrence across clinic visits that are not explained by the fixed temporal effects. *∈*_*patient*_ and *∈*_*visit*_ allow for residual pairwise correlations in co-occurrence that are not explained by the fixed, pairwise effects. Following the definition of the multivariate probit density, *∈*_*patient*_ and *∈*_*visit*_ are nested effects, such that the same *∈*_*patient*_ is added to to all of that patient’s visits, such that the variances of *∈*_*patient*_ and *∈*_*visit*_ must sum to one (i.e. *z ∼ N*(*µ, 1*)). These random effects are therefore structured as follows:

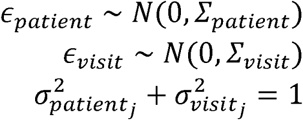

where *Σ*_*patient*_ and *Σ*_*visit*_ are *J*×*J* variance-covariance matrices, constrained so that the *j*^*th*^ variance parameters from the two matrices sum to one, for *j = 1*,…*J*. Therefore, 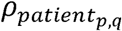 represents the pairwise correlation between HPV types that is measured among patients, which is derived from the variance-covariance matrix *Σ*_*patient*_. Then, 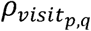 represents the pairwise correlation between HPV types that is measured between visits and within patients (i.e. longitudinally), which is derived from the variance-covariance matrix *Σ*_*visit*_.

We also model fixed effects of the time between visits (TBV) on persistence and colonization, to allow for the variability in when patients visited the clinic. The median TBV was 6.0 months with variance = 0.7 months, which we centered and scaled for use in the model. We allowed for fixed effects of TBV on the HPV type-specific probability of persisting 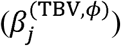 and the probability of colonizing 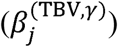. We hypothesized that the probability that an HPV type colonizes a patient increases with TBV, due to a longer period of risk, while the probability that a HPV type persists in the patient decreases with TBV, due to a longer time in which clearance may occur. The structure of these fixed effects is:

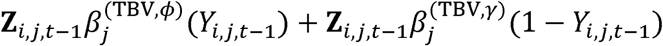

In this formula, ***Z*** is an *I* × *T* matrix that holds the centered and scaled values of TBV for each patient. This formula is added to *µ*_*ijt*_.

### Model inference

We coded our Bayesian model in *Stan* [51], an efficient, generalizable, statistical programming language, which employs adaptive Hamiltonian Monte Carlo (HMC) for model inference. We used hierarchical prior structures for the fixed effects and baseline prevalences, such that HPV type-level parameters were drawn from normal distributions with estimated means and standard deviations. Priors for mean values were all normally distributed with mean zero and standard deviation 1.5, which on the probit scale, allowed for a range between very weak and very strong effects. In other words, our priors were quite vague. Priors on standard deviations followed half-normal distributions centered at zero with standard deviation 1.0. For prior specification of the correlations in the random effects, we used the Cholesky LKJ correlation distribution. We made this distribution “vague” by setting the *η* term to 2, which “centers” the correlation matrix structure more towards an identity matrix (i.e. no correlation). Finally, we constrained the patient- and visit-level standard deviations to sum to one, to conform to the definition of the multivariate probit.

### Testing the model with synthetic data

We used synthetic data to verify that our model generates accurate inferences for data sets with various characteristics. In simulation 1, we tested whether our model could: (1) infer dynamics consistent with Simpson’s Paradox, meaning opposite correlations in among-patient effects versus among-visit effects, (2) infer dynamics given observations of rare species, and (3) infer weak inter-species interactions. We imposed these specific qualities because we suspected *a priori* that these characteristics could affect the model’s inference of HPV dynamics. We simulated data for a community of ten hypothetical pathogen strains sampled in 1500 patients, with each patient sampled 10 times. We assumed low but variable baseline probabilities of occurrence for each strain, matching the average baseline observed across the ten least prevalent HPV types in the HIM dataset. We further assumed positive patient-level correlations and negative observation-level correlations, such that correlations were equal across pathogen strain pairs 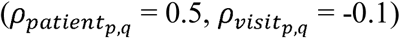. Pairwise effects on persistence and colonization 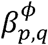 and 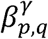 were drawn from normal distributions with mean zero and standard deviation 0.25.

In simulation 2, we sought to verify that large among-patient correlations (i.e. random effects) would not mask the inference of any fixed effects (even large ones), and that we could recover accurate parameter estimates with many fewer patients. We generated data for only 200 patients, each sampled 10 times. We also assumed a standard deviation of 1.35 among the fixed effects, leading to larger values of 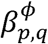 and 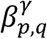 (i.e., much stronger effects on persistence and colonization). Finally, we assumed the among patient correlation was large 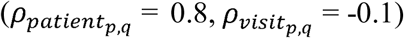.

In simulation 3, we tested whether our model-comparison approach (below) could reliably distinguish between the nested models for synthetic data. Specifically, we simulated data for a case in which fixed effects were present, but the random effect correlations were very low 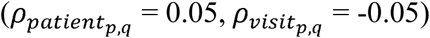. We then tested whether the model comparison would still show that the full model (with fixed and random effects) was still the best model, which would demonstrate that our model inference can accurately infer even weak correlations. All of our code for generating the synthetic data, as well as the data set itself, is available in our open-source repository: https://bitbucket.org/jrmihalj/hpv_jsdm.

### Fitting the model to the HIM data

Our first goal was to use our model to identify any interactions between HPV types that might warrant future epidemiological investigations. We therefore fit our full model and quantified the posterior distributions of the pairwise effects of HPV types on colonization and persistence rates. Our second goal was to understand the relative contributions of environmental effects, such as host-specific risk factors, and pairwise inter-type interactions to HPV community dynamics. We therefore fit four nested models of varying complexity. Model 1 has fixed, pairwise effects between HPV types, model 2 has residual correlations that account for environmental effects, and model 3, our full model, has both. Model 4 includes only baseline occurrence probabilities *α*_*j*_, and is therefore our null model. All of these models include the effects of the time between visits (TBV). We then compared the models’ out-of-sample predictive abilities using the leave-one-out information criterion (LOO-IC), estimated using Pareto-smoothed importance sampling in the *R* package “*loo*” [52]. Compared to the Watanabe-Akaike information criterion (WAIC), which is asymptotically equal to LOO-IC, the LOO-IC has been found to be more robust when using vague priors [53], as in our models. We considered two models to be substantially different if their LOO-IC values differed by 3, which is the common convention [54]. In practice, for a data set this large, small changes in overall goodness-of-fit could lead to very large changes in the likelihood when integrated across the many data points, and thus large differences in LOO-IC. We therefore emphasize that we use this model selection procedure as a heuristic to guide our understanding of community dynamics, rather than as a robust hypothesis test.

## Results

### Model validation with synthetic data

In simulation 1, our model accurately and precisely inferred dynamics from synthetic data consistent with Simpson’s Paradox, even when the data were sparse (Fig. 2). The model correctly inferred the low baseline probabilities of species occurrence (Fig. 2 A) and all patient-level correlations (Fig. 2 B). It also accurately estimated the majority of negative correlations at the observation level, although some inferred pairwise correlations were indistinguishable from zero (Fig. 2 C). This latter effect was not surprising, because we assumed a weak negative correlation (*ρ*_*visit*_ = -0.1). Importantly, although the model’s estimates of the magnitude of simulated correlations were sometimes incorrect, the model was unbiased with respect to the direction of the simulated correlations. The model also correctly estimated persistence 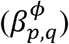 and colonization 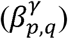, even when these effects were weak (Fig. 2 D,E). Finally, the model accurately recovered the effects of the time between visits on both persistence and colonization probabilities, which we assumed were the same for all pathogen strains (Fig. S2).

**Figure 2:**
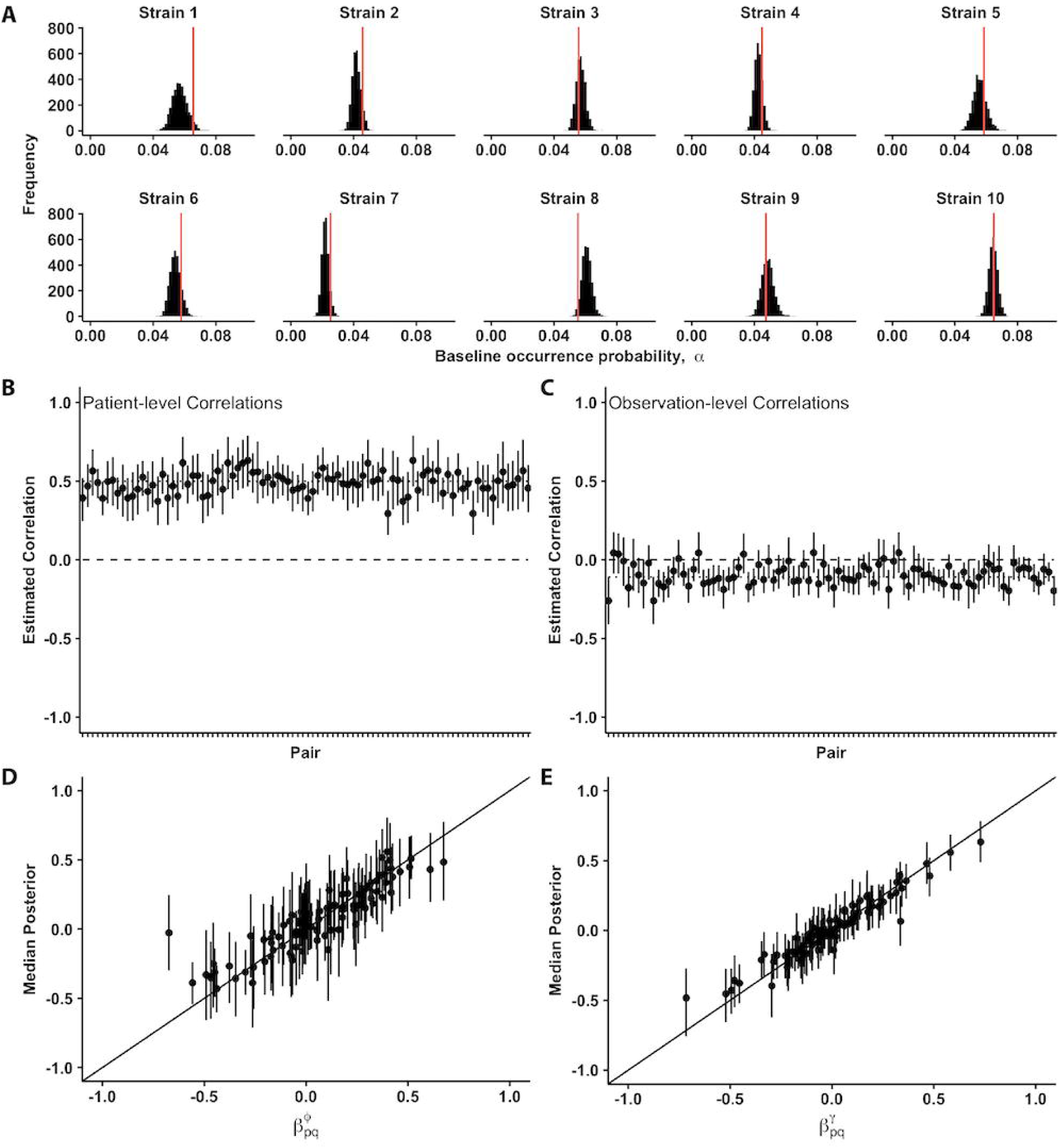
Inference of model parameters from synthetic data simulated for 10 pathogen strains across 1500 patients, where each patient was tested 10 times. **A** Recovery of baseline occurrence probability for the 10 strains. Red vertical line gives the true value. **B** Recovery of positive, pairwise correlations in among-patient random effects. Dashed line represents zero effect, while dotted line represents the true value (0.5). **C** Recovery of weakly negative, pairwise correlations in within-patient, observation-level random effects. Dashed line represents zero effect, while dotted line represents the true value (−0.11). **D** Recovery of fixed inter-strain effects on probability of strain persistence, with points representing median marginal posterior estimates and vertical lines representing 95% credible intervals (CI). The 1:1 relationship between true values and marginal posterior estimates of those values is also shown. Estimates falling on this line (within their 95% CI) represent accurate inferences. **E** Recovery of fixed inter-strain effects on probability of strain colonization, with 1:1 line as in panel D.

The model also performed well in simulations 2 and 3 with many fewer patients (200). In simulation 2, the model accurately inferred high among-patient correlations, and these high correlations did not bias the model’s inference of the fixed effects (Figs. S3, S4). In other words, the model can accurately distinguish between among-patient correlations and either strong or weak fixed effects on persistence and colonization. In simulation 3, the LOO-IC model selection approach showed that, even when we impose small patient- and visit-level correlations, the high accuracy of the model’s inference of these parameters leads to the full model (including both fixed and random effects) being chosen as the best model (Table S1, Figs. S5, S6).

### Metacommunity dynamics of HPV and model comparisons

In our full model, there were only a few interactions between HPV types that were worthy of future investigation, including several weakly negative effects on colonization probability (Fig. 3). Importantly, including these fixed effects and the random effects of patient-level and observation-level correlations led to a substantial improvement relative to a null model that accounts only for type-specific baseline occurrence probabilities, suggesting that the biology added to our model helps explain HPV community composition relative to the null model (Table 1). Based on LOO-IC selection, however, the most parsimonious model included only the random effects of patient-level and observation-level correlations, without pairwise interactions between the HPV types (Table 1). Pairwise inter-type interactions can thus be identified by our model, but the effect of these interactions is not strong enough to substantially mediate the overall community composition in this subset of 10 HPV types. The best model, which did not include these pairwise interactions, gives qualitatively similar insights for the random effects, meaning the patient-level and observation-level correlations, as our full model (Fig. S8).

**Table 1:** Comparison of candidate models using leave-one-out cross-validation. The table shows whether fixed and/or random effects were included (shown with a dot), the log-likelihood of the model fit (i.e. ℒ (*θ*|D)), and the LOO-IC. The standard error (SE) in the LOO-IC is shown to emphasize that the LOO-IC is an estimated statistic with error, but also that none of our LOO-IC values overlap within ± 2SE.

| HPV Interactions | Among-patient and Among-visit Correlations | Log-Likelihood | LOO-IC | SE LOO-IC |
| --- | --- | --- | --- | --- |
|  | • | -220310.1 | 708323.6 | 373.8 |
| • | • | -279574.9 | 825439.5 | 329.8 |
| • |  | -433036.9 | 1109717.0 | 465.3 |
|  |  | -432977.8 | 1130039.0 | 681.3 |

**Figure 3:**
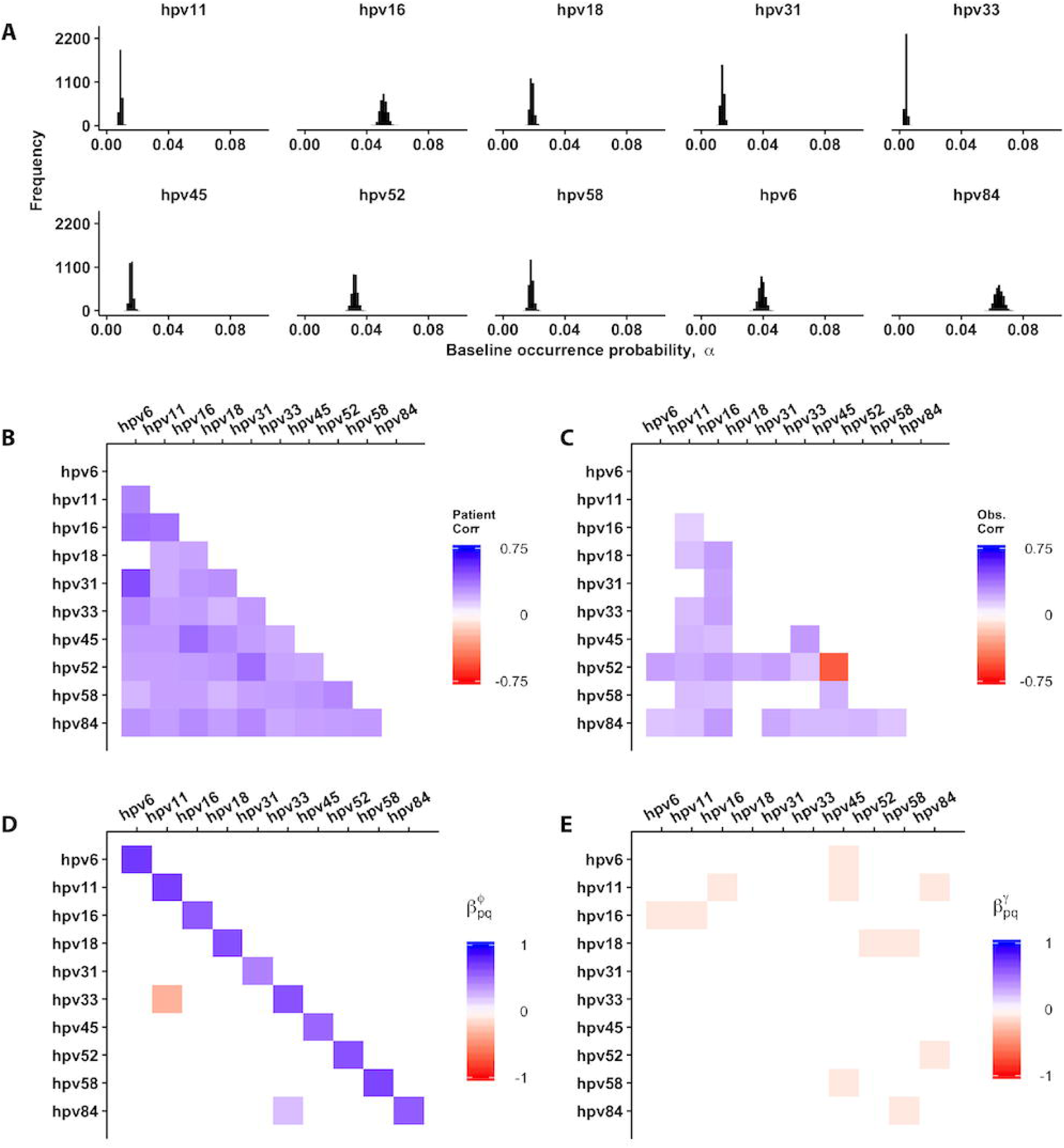
Inference of model parameters from the HIM data. **A** Estimate of the baseline occurrence probability for each HPV type. **B** Inferred correlations in among-patient random effects. **C** Inferred correlations in within-patient, observation-level random effects. **D** Inference of fixed inter-type effects on the probability of type persistence. **E** Inference of fixed inter-type effects on probability of type colonization.

The best model captured important qualitative aspects of the HPV dynamics, as well. The inferred baseline occurrence probability recovered the observed rank order of prevalence of the ten HPV types (Fig. 3A). The model confirmed that increasing values of TBV had positive effects on colonization probabilities 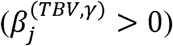 for all HPV types, but it had negative effects on persistence probabilities 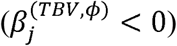 for all but two HPV types (Figs. S7, S8).

Patient-level correlations were positive for all but one pair of HPV types (Fig. 3B). These positive correlations suggest that there are shared environmental drivers across human hosts, in the form of risk factors. In the case of HPV52 and HPV58 (Fig. 3C), there are both positive patient-level and negative observation-level correlations. Positive observation-level correlations, or correlations within individuals over time, likely signal affinity for co-transmission, because in the models these effects are in addition to the pairwise effects on persistence and colonization. Negative observation-level correlations thus signal reduced affinity for co-transmission.

However, the negative observation-level correlations between HPV52 and HPV58 must be interpreted with caution, as they could reflect the masking of HPV58 detection by HPV52, a problem that has been documented in the linear array genotyping test used in the HIM study [43].

## Discussion

Our results suggest that HPV type coexistence is strongly driven by shared environmental characteristics. While the full model is able to estimate even sparse and weak (putative) interactions between HPV types, our model selection procedure suggests that these interactions are not important for explaining overall patterns of community turnover in HPV. The influence of patient-level correlations on HPV community dynamics suggests that HPV types segregate among hosts with shared traits. It is therefore likely that human subpopulations exist that could promote HPV type coexistence across space and time. This finding is consistent with epidemiological evidence of type-specific differences in the risk factors that promote HPV transmission [35, 55], and with another recent modeling study that characterized subtle differences in the profile of host-specific risk factors that affect infection with each type [56].

Model selection suggests that pairwise inter-type interactions that affect colonization and persistence probabilities do not strongly influence large-scale patterns of community turnover in this HPV data set. However, the full model identified several putative interactions worthy of future epidemiological investigations. In particular, it is possible that interactions could mediate the occurrence patterns of specific pairs of HPV types, even though model selection suggests that pairwise interaction effects have no meaningful effects on the HPV community dynamics as a whole. In other words, the community-level patterns could swamp out the patterns of specific HPV pairs. Further, by limiting our analysis to a subset of ten HPV types, it is possible that we by chance did not include HPV types that have larger effects on the community. Also, our model only estimates pairwise effects, and future studies could account for higher order interactions, which have been shown to be important in diverse competitive networks [57].

The results of our analysis complement the results of a previous, mechanistic model of HPV dynamics fitted to 6 HPV types of the HIM dataset [56]. The authors of this previous work formulated an epidemiological model that allowed for homologous immunity, a form of within-species competition, as well as the effects of 11 host-specific covariates. The best-fit version of this model included no homologous immunity for any of the six HPV types (HPV84, HPV62, HPV89, HPV16, HPV51, and HPV6), finding instead that previous infection with any type significantly increases the risk of re-infection with the same type. In our statistical model, this effect is further confirmed by the positive baseline persistence probabilities 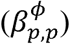 across all ten HPV types analyzed. In other words, all 10 HPV types had greater than 50% likelihood of persisting to the next clinic visit, which were on average 6 months apart, suggesting re-infection occurring in between visits. That study [56] also detected no pairwise interaction between two taxonomically similar types, HPV16 and HPV31, which had been hypothesized to compete through cross-immunity [58, 59]. Furthermore, the risk of initial infection with any HPV type was concentrated among high-risk subpopulations, which were linked to host-specific covariates. Taken together, the results of this previous analysis [56] suggest that both intra-specific and inter-specific competition are weak or absent in the HPV viral community, such that stabilizing competitive mechanisms cannot explain HPV diversity. Instead, diversity may depend on sustained infection within high-risk subpopulations specific to each HPV type. These findings are consistent with our finding that inter-type interactions have little effect on HPV community dynamics (Table 1). Furthermore, by showing how host-specific traits define niches that are used by different HPV types, the previous work [56] supports the importance of shared among-patient traits to explain patterns of co-occurrence.

While the different quantitative approaches between the previous study [56] and our study provide complementary results, there are important differences in the methods, applications, and conclusions. Ranjeva et al. [56] tested mechanistic biological models about type-specific HPV dynamics, whereas our approach allowed for the identification of statistical patterns in the community dynamics of multiple types. Also, our method can be generalized to any metacommunity that is sampled through time, rather than being specific to a pathogen community that interacts via cross-immunity, as modeled by Ranjeva et al. [56]. Indeed, our statistical framework is agnostic to the specific mechanisms of interactions. Instead our model specifies latent mechanisms that affect probabilities of persistence and colonization, which are estimated from the occurrence data.

We have shown that a relatively simple statistical model can be used to infer community dynamics, even in a system with rare species occurrences. Sparsity of observational data in real-world metacommunities generally limits the power of statistical models to correctly infer ecological effects [50, 60, 61]. We showed that our model can be used to infer opposing environmental and temporal dynamics from communities of rare species, and to detect weak interactions among rare species, which are the most common types of interactions in nature [62]. Inferring residual correlations with rare species requires a substantial amount of data, but, in the age of affordable, high-throughput sequencing technologies, such data can often be obtained easily. Moreover, our model accounts for the effects of unobserved environmental drivers, specifically host-specific risk-factors in the case of the HPV data, without having to specify covariates explicitly. This may be useful for analyzing large microbial communities, such as microbiome communities, in which the environmental drivers are unknown.

In classical joint-species distribution models, residual correlations in species occurrence are used to infer species interactions, but such residual correlations can arise instead from shared covariate responses that are not explicitly included in the model structure [2, 22]. Our model, however, does not rely on residual correlations to infer interspecies interactions *per se*. We use species occupancy at the previous time step to estimate lagged, pairwise effects of species’ occurrences on the probabilities of persistence and colonization of cohabitating species. Residual correlations in our models instead account for latent environmental covariates, such as unmeasured host-specific traits. Although our statistical modeling approach can thus identify important signatures of species interactions, mechanistic models and experimentation are nevertheless required to rigorously test hypotheses about species interactions. Furthermore, we estimate interspecies effects on persistence and colonization using a one-timestep lag, which requires that the timescale of the species interactions be equal to the timescale of observations. This assumption may not always hold. Our method is therefore best used to refine testable hypotheses from observed dynamics of large community assemblages, such as microbiome assemblages, in a computationally-feasible manner, rather than as a final step in inferring interactions.

A final caveat is that our models do not allow for dynamics that occur between observations. Given two consecutive observations of a species, our models instead assume that there is either persistence over the entire interval, or that at most one extinction or colonization has occurred. This assumption may result in bias in communities that are poorly sampled relative to the timescale of the dynamics. Indeed, recent evidence shows that standard joint-species distribution modeling approaches cannot accurately capture simulated predator-prey dynamics, especially if habitats are relatively homogeneous, probably because of non-linear dynamics [2]. This problem is likely to be important for non-linear host-pathogen dynamics as well, and should be a subject of future simulation efforts. Our dataset however spans a wide diversity of patients, and includes the effects of the time between visits, which should limit this type of bias.

## Competing Interests

The authors declare no competing financial interests

## References

1. Gotelli NJ, Graves GR. Null Models in Ecology. 1996. Smithsonian Institution Press, Washington, D.C.

2. Zurell D, Pollock LJ, Thuiller W. Do joint species distribution models reliably detect interspecific interactions from co-occurrence data in homogenous environments? Ecography (Cop).

3. Keeling M, Rohani P. Modeling infectious diseases in humans and animals. 2008. Princeton University Press.

4. Godsoe W, Franklin J, Blanchet FG. Effects of biotic interactions on modeled species’ distribution can be masked by environmental gradients. Ecol Evol 2017; 7: 654–664.

5. Faust K, Raes J. Microbial interactions: From networks to models. Nat Rev Microbiol 2012; 10: 538–550.

6. Seabloom EW, Borer ET, Gross K, Kendig AE, Lacroix C, Mitchell CE, et al. The community ecology of pathogens: coinfection, coexistence and community composition. Ecol Lett 2015; 18: 401–415.

7. Diamond JM. Assembly of species communities. Ecol Evol communities 1975; 342–444.

8. Connor EF, Simberloff D. The Assembly of Species Communities: Chance or Competition? Ecology 1979; 60: 1132.

9. Gotelli NJ, McCabe DJ. Species cooccurrence: A meta-analysis of JM Diamond’s assembly rules model. Ecology 2002; 83: 2091–2096.

10. Fisher CK, Mehta P. Identifying keystone species in the human gut microbiome from metagenomic timeseries using sparse linear regression. PLoS One 2014; 9: 1–10.

11. Weiss S, Van Treuren W, Lozupone C, Faust K, Friedman J, Deng Y, et al. Correlation detection strategies in microbial data sets vary widely in sensitivity and precision. ISME J 2016; 10: 1669–1681.

12. Ruan Q, Dutta D, Schwalbach MS, Steele JA, Fuhrman JA, Sun F. Local similarity analysis reveals unique associations among marine bacterioplankton species and environmental factors. Bioinformatics 2006; 22: 2532–2538.

13. Xia LC, Ai D, Cram JA, Liang X, Fuhrman JA, Sun F. Statistical significance approximation in local trend analysis of high-throughput time-series data using the theory of Markov chains. BMC Bioinformatics 2015; 16: 1–14.

14. Stein RR, Bucci V, Toussaint NC, Buffie CG, Rätsch G, Pamer EG, et al. Ecological Modeling from Time-Series Inference: Insight into Dynamics and Stability of Intestinal Microbiota. PLoS Comput Biol 2013; 9: 31–36.

15. Carrara F, Giometto A, Seymour M, Rinaldo A, Altermatt F. Experimental evidence for strong stabilizing forces at high functional diversity of microbial communities. Press 2015; 96: 1340–1350.

16. Cardona C, Weisenhorn P, Henry C, Gilbert JA. Network-based metabolic analysis and microbial community modeling. Curr Opin Microbiol 2016; 31: 124–131.

17. Coenen AR, Weitz JS. Limitations of Correlation-Based Inference in Complex Virus-Microbe Communities. mSystems 2018; 3: 7–9.

18. Ovaskainen O, Hottola J, Siitonen J. Modeling species co-occurrence by multivariate logistic regression generates new hypotheses on fungal interactions. Ecology 2010; 91: 2514–2521.

19. Ovaskainen O, Tikhonov G, Norberg A, Blanchet FG, Duan L, Dunson D, et al. How to make more out of community data? A conceptual framework and its implementation as models and software. Ecol Lett 2017; 20: 561–576.

20. Ovaskainen O, Abrego N, Halme P, Dunson D. Using latent variable models to identify large networks of species-to-species associations at different spatial scales. Methods Ecol Evol 2015.

21. Sebastián-González E, Sánchez-Zapata JA, Botella F, Ovaskainen O. Testing the heterospecific attraction hypothesis with time-series data on species co-occurrence. Proc R Soc B Biol Sci 2010; 277: 2983–2990.

22. Pollock LJ, Tingley R, Morris WK, Golding N, O’Hara RB, Parris KM, et al. Understanding co-occurrence by modelling species simultaneously with a Joint Species Distribution Model (JSDM). Methods Ecol Evol 2014; 5: 397–406.

23. Joseph MB, Preston DL, Johnson PTJ. Integrating occupancy models and structural equation models to understand species occurrence. Ecology 2016; 97: 765–775.

24. Ma Y, Madupu R, Karaoz U, Nossa CW, Yang L, Yooseph S, et al. Human papillomavirus community in healthy persons, defined by metagenomics analysis of human microbiome project shotgun sequencing data sets. J Virol 2014; 88: 4786–4797.

25. Frazer IH. Interaction of human papillomaviruses with the host immune system: A well evolved relationship. Virology 2009; 384: 410–414.

26. Todd RW, Roberts S, Mann CH, Luesley DM, Gallimore PH, Steele JC. Human papillomavirus (HPV) type 16-specific CD8+ T cell responses in women with high grade vulvar intraepithelial neoplasia. Int J Cancer 2004; 108: 857–862.

27. Dunne EF, Unger ER, Mcquillan G, Swan DC, Patel SS, Markowitz LE. Prevalence of HPV Infection. 2014; 297: 813–819.

28. Markowitz LE, Hariri S, Lin C, Dunne EF, Steinau M, McQuillan G, et al. Reduction in human papillomavirus (HPV) prevalence among young women following HPV vaccine introduction in the United States, National Health and Nutrition Examination Surveys, 2003-2010. J Infect Dis 2013; 208: 385–393.

29. Joura EA, Giuliano AR, Iversen O-E, Bouchard C, Mao C, Mehlsen J, et al. A 9-Valent HPV Vaccine against Infection and Intraepithelial Neoplasia in Women. N Engl J Med 2015; 372: 711–723.

30. Bernard E, Pons-Salort M, Favre M, Heard I, Delarocque-Astagneau E, Guillemot D, et al. Comparing human papillomavirus prevalences in women with normal cytology or invasive cervical cancer to rank genotypes according to their oncogenic potential: a metaanalysis of observational studies. BMC Infect Dis 2013; 13: 373.

31. Durham DP, Poolman EM, Ibuka Y, Townsend JP, Galvani AP. Reevaluation of epidemiological data demonstrates that it is consistent with cross-immunity among human papillomavirus types. J Infect Dis 2012; 206: 1291–1298.

32. Bernard H-U, Burk RD, Chen Z, van Doorslaer K, Hausen H zur, de Villiers E-M. Classification of papillomaviruses (PVs) based on 189 PV types and proposal of taxonomic amendments. Virology 2010; 401: 70–79.

33. Giuliano AR, Lazcano-Ponce E, Villa LL, Flores R, Salmeron J, Lee J-H, et al. The human papillomavirus infection in men study: human papillomavirus prevalence and type distribution among men residing in Brazil, Mexico, and the United States. Cancer Epidemiol Biomarkers Prev 2008; 17: 2036–2043.

34. Giuliano AR, Lazcano E, Villa LL, Flores R, Salmeron J, Lee J-H, et al. Circumcision and sexual behavior: factors independently associated with human papillomavirus detection among men in the HIM study. Int J Cancer 2009; 124: 1251–1257.

35. Nyitray AG, Carvalho RJ, Baggio ML, Beibei L, Abrahamsen M, Papenfuss M, et al. Age-Specific Prevalence of and Risk Factors for Anal Human Papillomavirus (HPV) among Men Who Have Sex with Women and Men Who Have Sex with Men_J: The HPV in Men (HIM) Study. J Infect Dis 2011; 203: 49–57.

36. Nyitray AG, Carvalho Da Silva RJ, Baggio ML, Smith D, Abrahamsen M, Papenfuss M, et al. Six-month incidence, persistence, and factors associated with persistence of anal human papillomavirus in men: The HPV in men study. J Infect Dis 2011; 204: 1711–1722.

37. Kahn J a, Brown DR, Ding L, Widdice LE, Shew ML, Glynn S, et al. Vaccine-type human papillomavirus and evidence of herd protection after vaccine introduction. Pediatrics 2012; 130: 249–256.

38. Wheeler CM, Castellsague X, Garland SM, Szarewski A, Paavonen J, Naud P, et al. Cross-protective efficacy of HPV-16/18 AS04-adjuvanted vaccine against cervical infection and precancer caused by non-vaccine oncogenic HPV types: 4-year end-of-study analysis of the randomised, double-blind PATRICIA trial. Lancet Oncol 2012; 13: 100–110.

39. Rousseau MC, Pereira JS, Prado JC, Villa LL, Rohan TE, Franco EL. Cervical coinfection with human papillomavirus (HPV) types as a predictor of acquisition and persistence of HPV infection. J Infect Dis 2001; 184: 1508–1517.

40. Liaw KL. A prospective study of human papillomavirus (HPV) type 16 DNA detection by polymerase chain reaction and its association with acquisition and persistence of other HPV types. J Infect Dis 2001; 183: 8–15.

41. Chaturvedi AK, Myers L, Hammons AF, Clark RA, Dunlap K, Kissinger PJ, et al. Prevalence and clustering patterns of human papillomavirus genotypes in multiple infections. Cancer Epidemiol Biomarkers Prev 2005; 14: 2439–2445.

42. Chaturvedi AK, Katki HA, Hildesheim A, Rodríguez AC, Quint W, Schiffman M, et al. Human papillomavirus infection with multiple types: Pattern of coinfection and risk of cervical disease. J Infect Dis 2011; 203: 910–920.

43. Tota JE, Ramanakumar A V, Jiang M, Dillner J, Walter SD, Kaufman JS, et al. Epidemiologic approaches to evaluating the potential for human papillomavirus type replacement postvaccination. Am J Epidemiol 2013; 178: 625–634.

44. Elbasha EH, Galvani AP. Vaccination against multiple HPV types. Math Biosci 2005; 197: 88–117.

45. Murall CL, McCann KS, Bauch CT. Revising ecological assumptions about Human papillomavirus interactions and type replacement. J Theor Biol 2014; 350: 98–109.

46. Poolman EM, Elbasha EH, Galvani AP. Vaccination and the evolutionary ecology of human papillomavirus. Vaccine 2008; 26: 25–30.

47. Giuliano AR, Lee J-H, Fulp W, Villa LL, Lazcano E, Papenfuss MR, et al. Incidence and clearance of genital human papillomavirus infection in men (HIM): a cohort study. Lancet 2011; 377: 932–940.

48. Han JJ, Beltran TH, Song JW, Klaric J, Choi YS. Prevalence of Genital Human Papillomavirus Infection and Human Papillomavirus Vaccination Rates Among US Adult Men. JAMA Oncol 2017; 3: 810–816.

49. Giuliano AR, Nyitray AG, Kreimer AR, Pierce Campbell CM, Goodman MT, Sudenga SL, et al. EUROGIN 2014 roadmap: Differences in human papillomavirus infection natural history, transmission and human papillomavirus-related cancer incidence by gender and anatomic site of infection. Int J Cancer 2015; 136: 2752–2760.

50. Dorazio RM, Kéry M, Royle JA, Plattner M. Models for inference in dynamic metacommunity systems. Ecology 2010; 91: 2466–2475.

51. Carpenter B, Gelman A, Hoffman MD, Lee D, Goodrich B, Betancourt M, et al. *Stan*_J: A Probabilistic Programming Language. J Stat Softw 2017; 76.

52. Vehtari A, Gelman A, Gabry J. loo: Efficient leave-one-out cross-validation and WAIC for Bayesian models. 2016.

53. Vehtari A, Gelman A, Gabry J. Practical Bayesian model evaluation using leave-one-out cross-validation and WAIC. Stat Comput 2016; 27: 1–20.

54. Gelman A, Carlin JB, Stern HS, Rubin DB. Bayesian Data Analysis, 2nd ed. 2003. Chapman & Hall/CRC.

55. Albero G, Castellsagué X, Lin H-Y, Fulp W, Villa LL, Lazcano-Ponce E, et al. Male circumcision and the incidence and clearance of genital human papillomavirus (HPV) infection in men: the HPV Infection in men (HIM) cohort study. BMC Infect Dis 2014; 14: 75.

56. Ranjeva SL, Baskerville EB, Dukic V, Villa LL, Lazcano-Ponce E, Giuliano AR, et al. Recurring infection with ecologically distinct HPV types can explain high prevalence and diversity. Proc Natl Acad Sci U S A 2017; 114: 13573–13578.

57. Levine JM, Bascompte J, Adler PB, Allesina S. Beyond pairwise mechanisms of species coexistence in complex communities. Nature 2017; 546: 56–64.

58. Draper E, Bissett SL, Howell-Jones R, Edwards D. Neutralization of non-vaccine human papillomavirus pseudoviruses from the A7 and A9 species groups by bivalent HPV vaccine sera. Vaccine 2011; 29: 8585–8590.

59. Kemp TJ, Hildesheim A, Safaeian M, Dauner JG, Pan Y, Porras C, et al. HPV16/18 L1 VLP vaccine induces cross-neutralizing antibodies that may mediate cross-protection. Vaccine 2011; 29: 2011–2014.

60. Mihaljevic JR, Joseph MB, Johnson PT. Using multispecies occupancy models to improve the characterization and understanding of metacommunity structure. Ecology 2015; 96: 1783–1792.

61. Warton DI, Blanchet FG, O’Hara RB, Ovaskainen O, Taskinen S, Walker SC, et al. So Many Variables: Joint Modeling in Community Ecology. Trends Ecol Evol. 2015. Elsevier Ltd., 30: 766–779

62. Chesson P. Mechanisms of Maintenance of Species Diversity. Annu Rev Ecol Syst 2000; 31: 343–366.

