## Supporting Information for "Untangling the dynamics of persistence and colonization in microbial communities"

### Supporting Information (SI)

#### *Ranjeva et al. 2019, The ISME Journal*

#### ADDITIONAL METHODS

#### Subset of HIM data included in the analysis

We excluded individuals that failed to meet the full eligibility criteria described by the HIM study. The criteria included: ages 18 to 70 years; residents of one of three sites – Sao Paulo, Brazil; Morelos, Mexico; or southern Florida, United States; no prior diagnosis of penile or anal cancers; no prior history of genital or anal warts; no symptoms of a sexually transmitted infection at baseline or recent treatment for a sexually transmitted infection; no history of participation in an HPV vaccine study; and no history of HIV or AIDS. We identified 3,656 eligible participants from the 4,123 men enrolled in the HIM study as of October 2014. For each of the 10 HPV types that we analyzed, we include in our data the binary infection status of each man at each clinic visit. We also include the length of time between consecutive clinic visits.

#### Type-specific HPV prevalence over follow-up

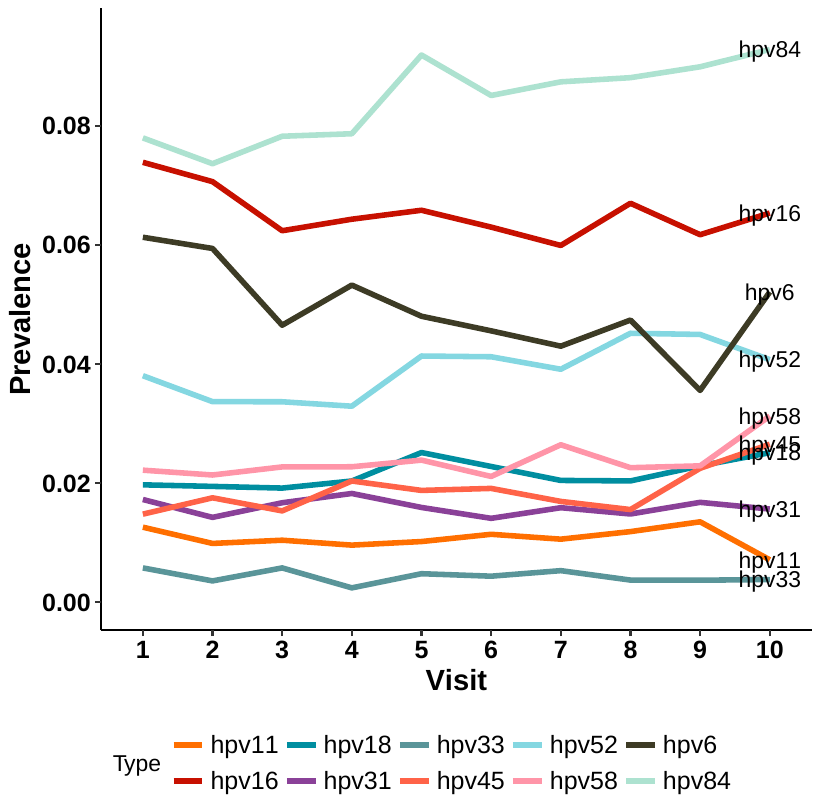
We calculated the prevalence of the 10 HPV types included in the analysis at each visit (Fig. S1). Note that, because individuals varied in their visit dates, the prevalence at each visit is a time-averaged estimate. The data show that the expected distribution of HPV types in the metacommunity is consistent across visits.

**Figure S1**. Observed visit-level prevalence of each of the 10 HPV types included in this analysis.

#### Stan model details

All of our code to run the Stan model is provided in our open-source repository <https://bitbucket.org/jrmihalj>, but we will briefly describe the fitting routine here. For each nested model, we ran three MCMC chains in parallel on the high performance computing (HPC) clusters at the University of Chicago (Center for Research Informatics) or at Northern Arizona University (. Each chain ran for 5000 iterations with a 2000 iteration warm-up period, and we thinned the samples by three, giving us a total of 1000 posterior samples from each chain. Parameter samples were stored as tables in an SQLite database for later processing. Due to the large number of columns of the log-likelihood table, we split this table into sub-components before storage. We monitored convergence with the Gelman-Rubin ($\hat{R}$) statistic, and we conducted several standard visual diagnostics to check MCMC chain performance. All models converged after 5000 iterations, and no problems were observed in the MCMC chains.

#### ADDITIONAL SIMULATION RESULTS

#### Simulation 1

#### *Time between visits - Simulation 1*

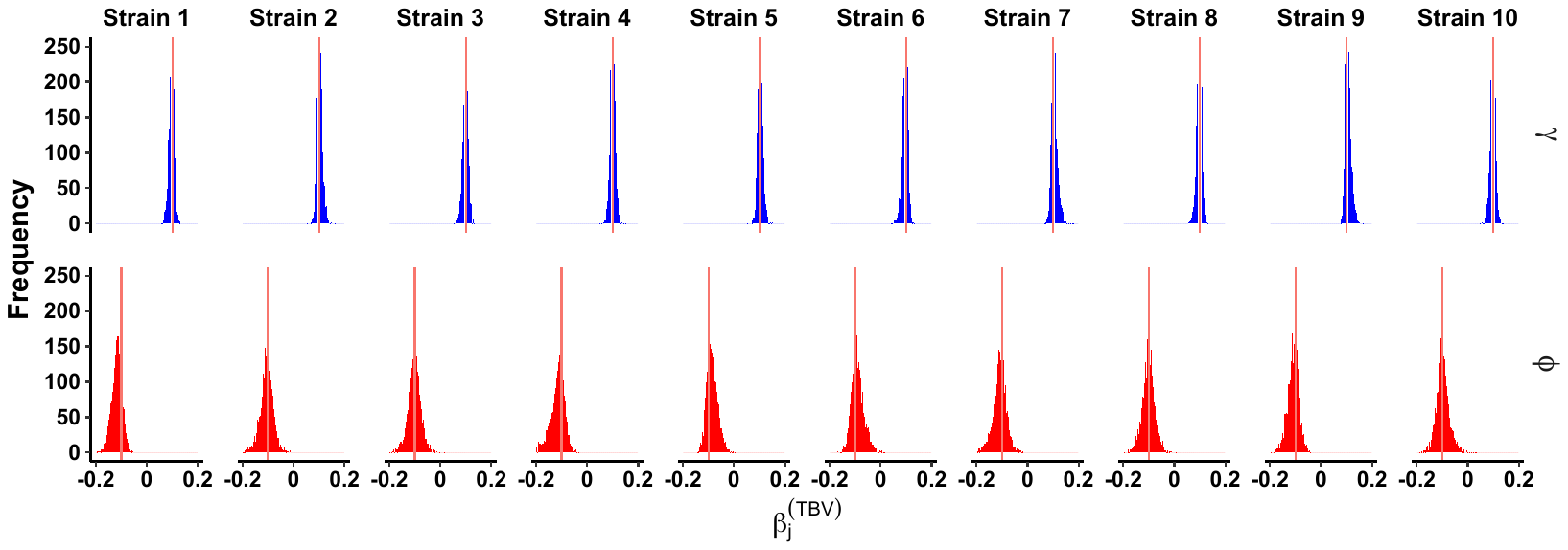
**Figure S2:** Effects of time between visit (TBV) on colonization (top row) and persistence (bottom row) probabilities for each of 10 simulated pathogen strains, from simulation 1. These results are generated from the full model, which has both fixed effects of pairwise interactions, as well as patient-level and observation-level correlations among residuals. Blue histograms are effects greater than zero, while red histograms are effects less than zero, based on a 95% credibal intervals (CI) that does not overlap zero. The true, simulated values are shown as red vertical lines.

#### Simulation 2

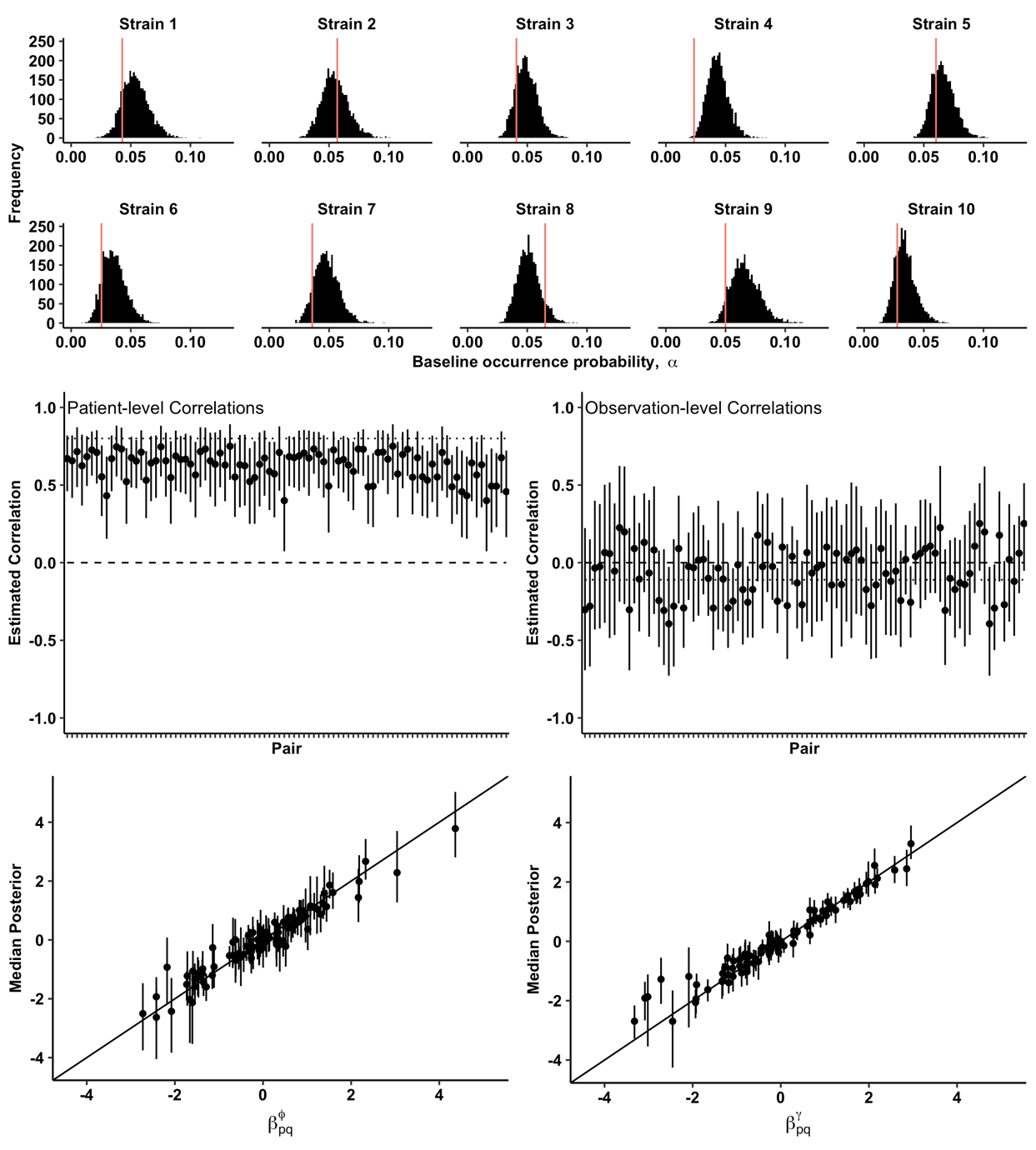
In this simulation, we generated data for only 200 patients, each sampled 10 times. We also assumed a standard deviation of 1.35 among the fixed effects, leading to larger values of $\beta_{p,q}^{\phi}$ and $\beta_{p,q}^{\gamma}$ (i.e., much stronger effects on persistence and colonization). Finally, we assumed the among patient correlation was large ($\rho_{patient_{p,q}}$ = 0.8,$\rho_{visit_{p,q}}$ = -0.1). Figure S3 shows that we accurately infer model parameters from this synthetic data set.

**Figure S3:** Inference of model parameters from synthetic data simulated for 10 pathogen strains across 200 patients, where each patient was tested 10 times, with large variance in fixed effects (leading to strong positive and negative fixed effects), and with large patient-level correlations. Figure legend same as Figure 2 of the main text.

#### *Time between visits – Simulation 2*

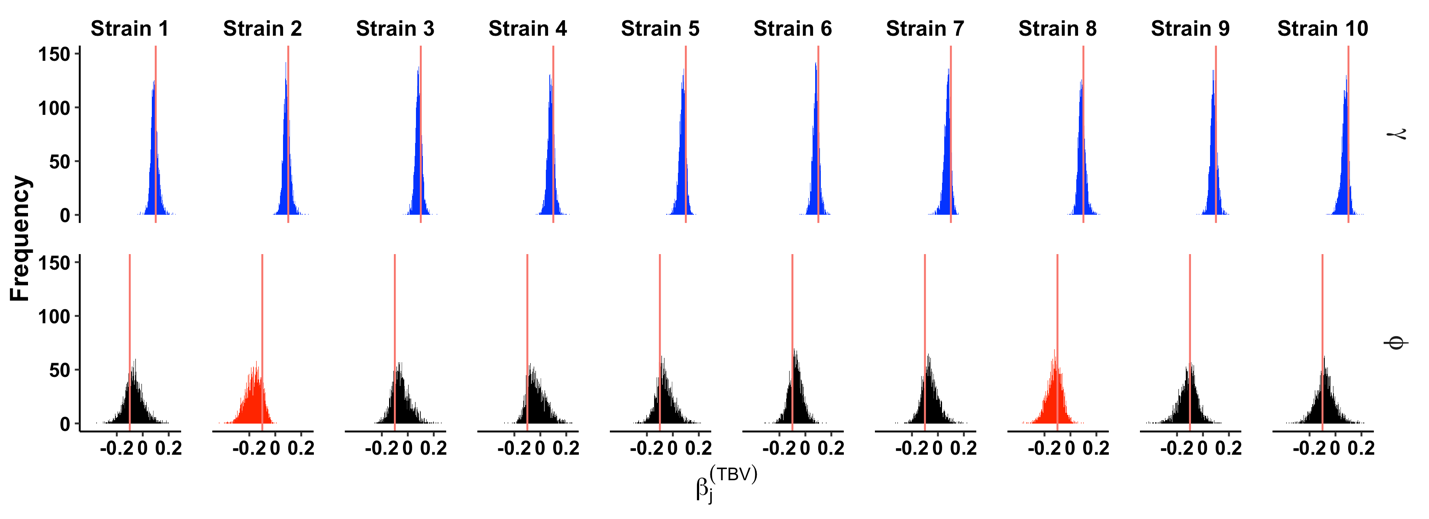

#### Figure S4: Effects of time between visit (TBV) on colonization (top row) and persistence (bottom row) probabilities for each strain type in simulation 2. Legend follows Figure S2.

#### Simulation 3

#### In simulation 3, we tested whether our model-comparison approach (below) could reliably distinguish between the nested models for synthetic data. Specifically, we simulated data for a case in which fixed effects were present, but the random effect correlations were very low ($\boldsymbol{\rho}_{\boldsymbol{patien}\boldsymbol{t}_{\boldsymbol{p,q}}}$ = 0.05,$\boldsymbol{\rho}_{\boldsymbol{visi}\boldsymbol{t}_{\boldsymbol{p,q}}}$ = -0.05). We then tested whether the model comparison would still show that the full model (with fixed and random effects) was still the best model, which would demonstrate that our model inference can accurately infer even weak correlations.

#### We accurately infer the parameters from this synthetic data set (Fig S5). Furthermore, model selection based on LOO-IC shows that the full model (with both fixed and random effects) was the best model. Thus, the model inference is precise enough to capture even small correlations among patients or between visits.

| HPV Interactions | Among-patient and Among-visit Correlations | Log-Likelihood | LOO-IC | SE LOO-IC |
| --- | --- | --- | --- | --- |
| • | • | **-24575.7** | **49151.4** | **111.5** |
|  | • | -26137.9 | 52275.9 | 125.9 |
|  |  | -36136.42 | 72272.85 | 139.30 |
| • |  | -38449.50 | 76898.99 | 135.04 |

**Table S1:** LOO-IC model selection for model fits to data in simulation 3. Legend as in Table 1 of the main text. The best model is bolded.

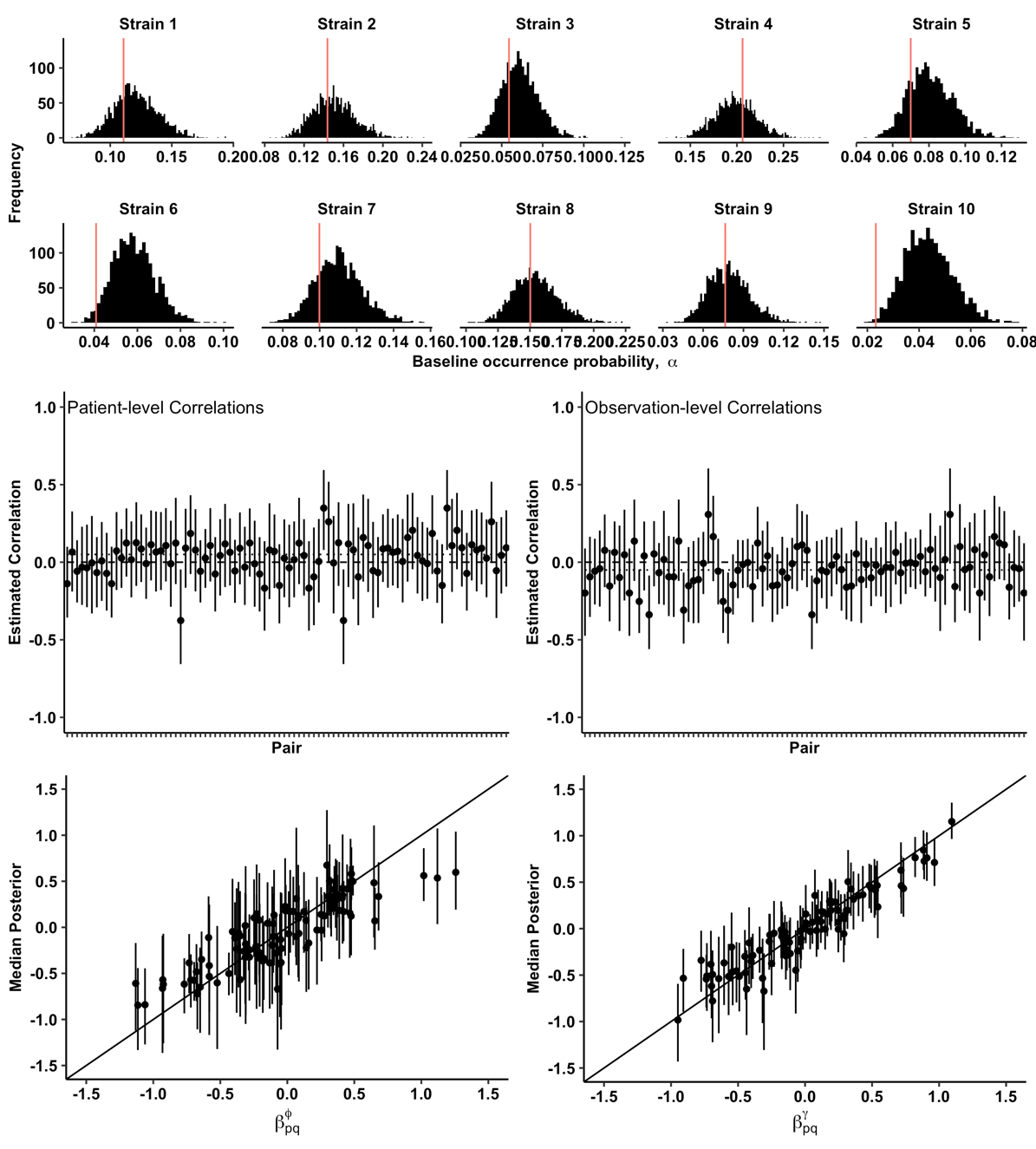

**Figure S5:** Inference of model parameters from synthetic data simulated for 10 pathogen strains across 200 patients, where each patient was tested 10 times, with small patient-level correlations. Figure legend same as Figure 2 of the main text.

*Time between visits – Simulation 3*

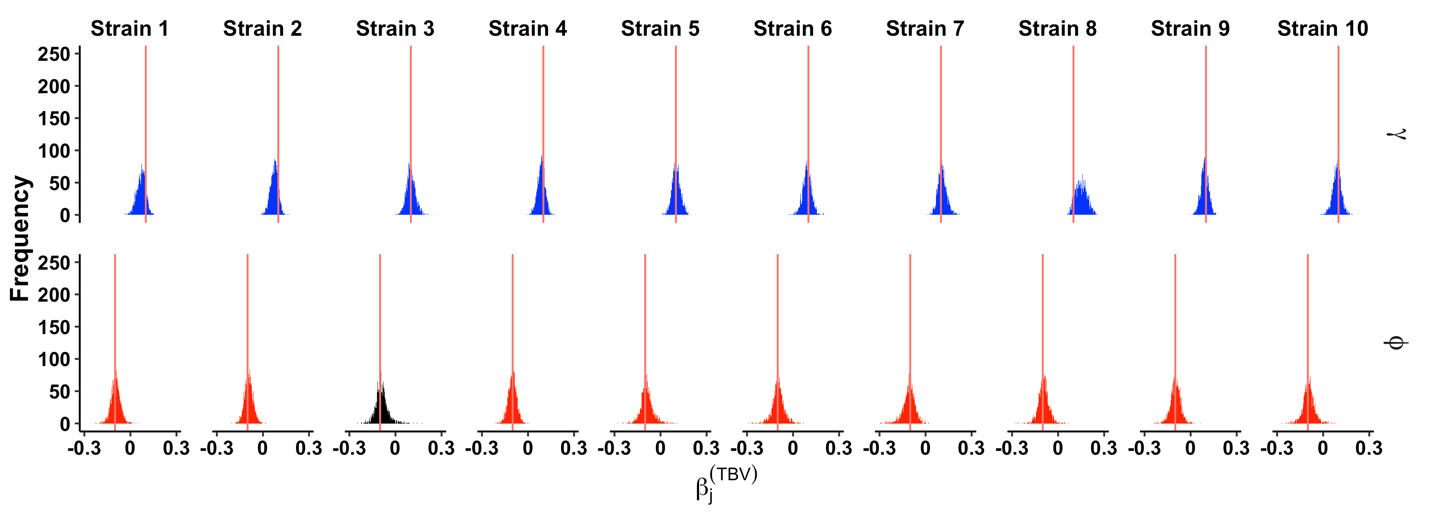

#### Figure S6: Effects of time between visit (TBV) on colonization (top row) and persistence (bottom row) probabilities for each strain type in simulation 3. Legend follows Figure S2.

#### ADDITIONAL RESULTS FROM HPV MODEL-FITTING

#### Time between visits - HPV dataset

Here we display the effects of time between visit (TBV) on persistence and colonization probabilities for the HIM dataset, using the full model that includes both correlations and fixed, pairwise interactions (Fig. S7).

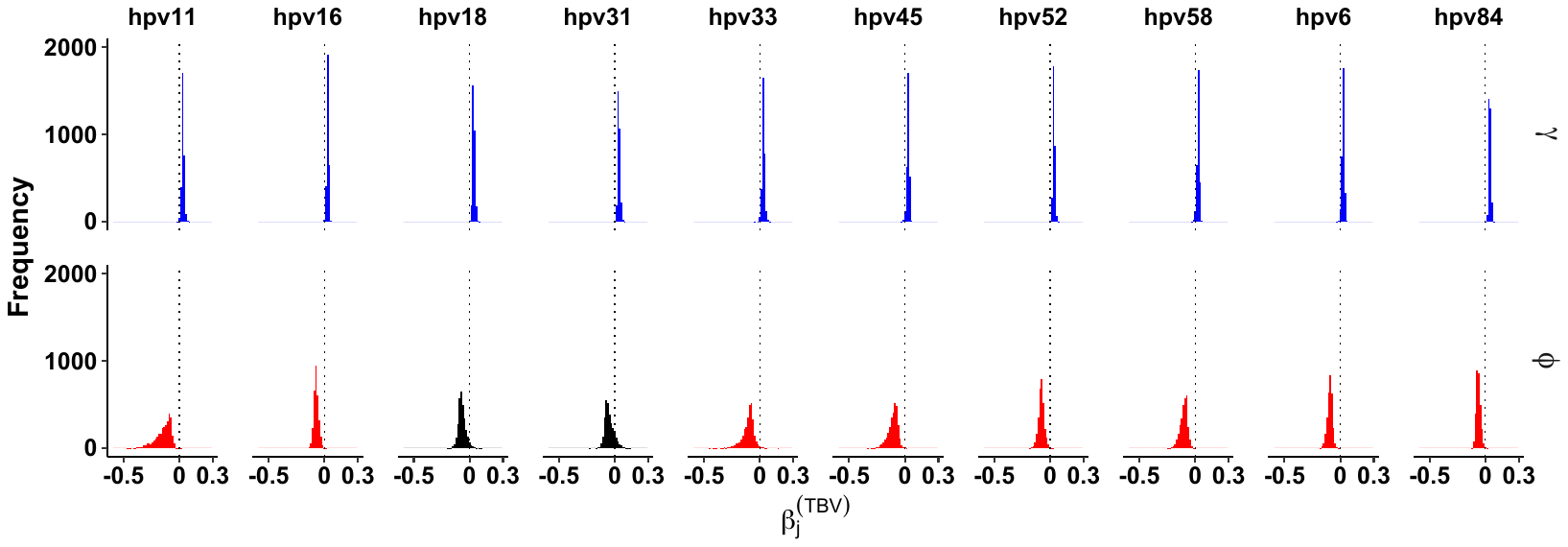

#### Figure S7: Effects of time between visit (TBV) on colonization (top row) and persistence (bottom row) probabilities for each HPV type in the HIM data set. Legend follows Figure S2.

#### Results from “best” model, with no pairwise interaction effects

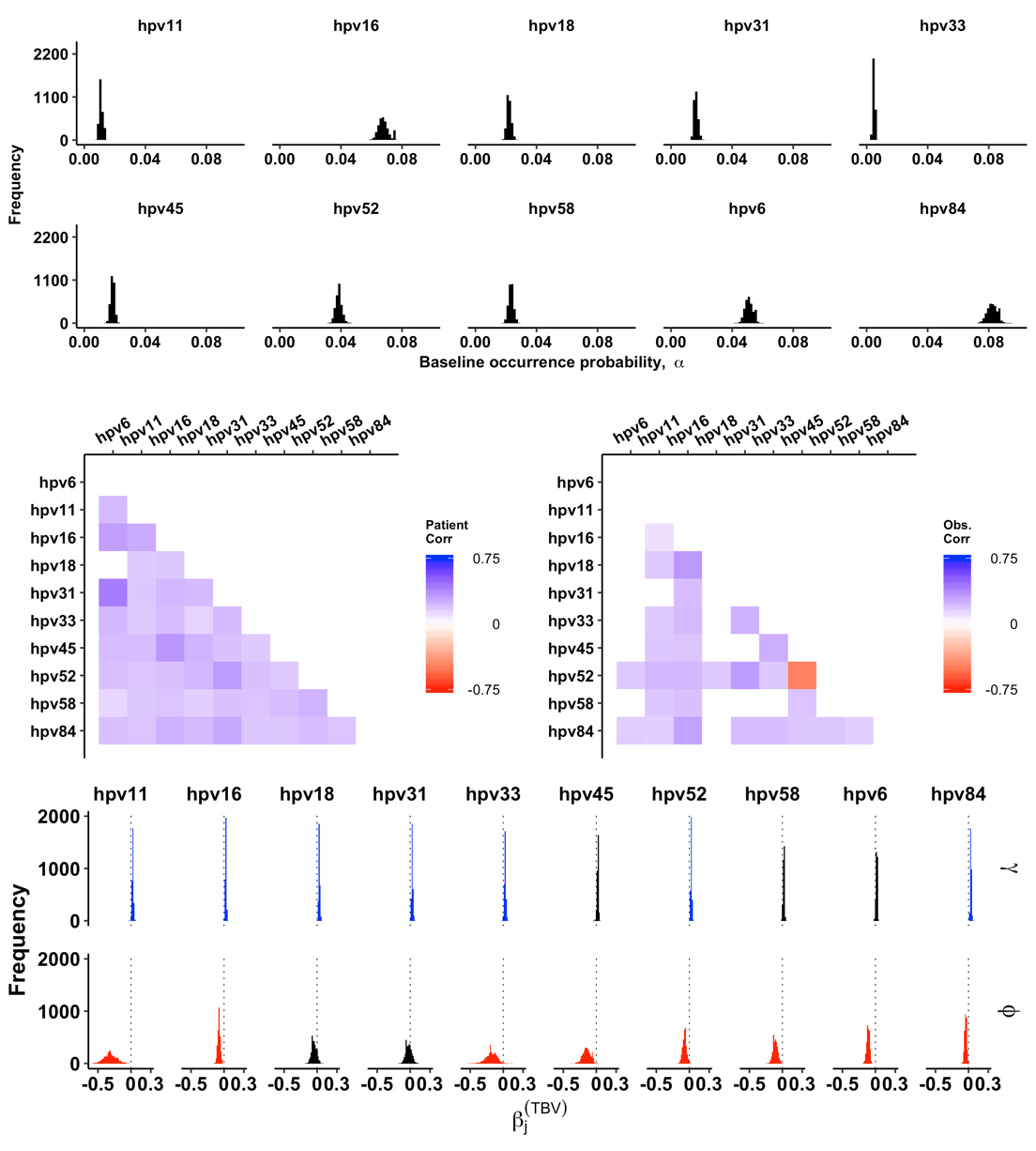
The figure below displays the results from the most preferred model, which includes the random effects (i.e. patient-level and observation-level correlations among HPV types), but does not include pairwise effects on persistence and colonization probabilities (Fig. S8).

**Figure S8**: Inference of model parameters from the HIM data, using the “best” model, which only has patient-level and observation-level correlations among types, but does not have fixed effects on persistence and colonization.
